# Learnable Graph Network Model (LGNM): A Physics Constrained Graph Neural Network with Quantum Hamiltonian Learning

**DOI:** 10.64898/2026.07.14.738428

**Authors:** Bhuwan Sharma, Chiranjib Sarkar

## Abstract

Elastic Network Models (ENMs), particularly the Gaussian Network Model (GNM) and its distance-weighted variant (mENM), predict per-residue protein flexibility from *C*_*α*_ contact graphs at low computational cost. Their central limitation is the assumption of uniform spring constants, which ignores the chemical identity, burial depth, and evolutionary conservation of individual residue contacts. We introduce the Learnable Graph Network Model (LGNM), a heterogeneous ENM in which per-edge spring constants *θ*_*ij*_ = *f*_*i*_ · *f*_*j*_ · (*d*_*c*_*/r*_*ij*_)^2^ are parameterised by per-residue flexibility coefficients {*f*_*i*_} predicted by a physics-constrained Graph Neural Network (GNN). The GNN is trained on molecular dynamics (MD)-derived root-mean-square fluctuation (RMSF) profiles from 413 proteins in the ATLAS database, using fold-disjoint CATH superfamily splits. The learning objective is an instance of the Quantum Neural PDE (QNPDE) Hamiltonian learning framework, with *K* = 3 operator types enabling an *O*(*K*) quantum gradient versus *O* (*N*^3^) classical pseudo-inversion. On 91 held-out test proteins, LGNM achieves mean per-protein Pearson correlation *r* = 0.8549± 0.1055, versus *r* = 0.8024 ± 0.1167 for mENM (Δ*r* = +0.0525; 77/91 proteins improved). The implementation of this methedology is aviliable at https://lgnm.compbiosysnbu.in/ allowing researchers to evaluate flexibility and downstream processses.

## 1 Introduction

Protein flexibility governs essential biological processes, including enzyme catalysis, allosteric signalling, molecular recognition, and intrinsic protein disorder [1; 2; 3]. Accurately predicting residues which are rigid and flexible from atomic structure would enable rapid screening of drug targets, designing of stable enzymes, and mechanistic interpretation of disease mutations—at a fraction of the cost of all-atom molecular dynamics (MD) simulation.

Elastic Network Models (ENMs) [4; 5] address this need by representing a protein as a network of harmonic springs between *C*_*α*_ atoms within a cutoff distance. The Gaussian Network Model (GNM) [4] assigns a single uniform spring constant *γ* = 1 to every contact edge; per-residue fluctuations are obtained analytically from the pseudo-inverse of the Kirchhoff (graph Laplacian) matrix. The Anisotropic Network Model (ANM) [5] extends GNM to a 3*N* × 3*N* Hessian capturing directional information. Distance-weighted modifications (mENM) set *γ*_*ij*_ = (*d*_*c*_*/r*_*ij*_)^*p*^ [6], replacing the binary cutoff by scaling the spring constants inversely with the distance between residues. While models likeFitNMA [7] go further by assigning per-atom flexibility constants {*k*_*i*_} fitted by nonlinear optimisation to crystallographic B-factors. Despite these architectural variations, existing models suffer from a fundamental trade-off: they either assume rigid, physically uniform springs, or rely on optimization techniques that fit per-atom flexibility constants to static crystallographic B-factors. These fitting approaches are inherently limited because they require slow, per-protein nonlinear optimization, difficulty to generalize to larger or unseen structures. Furthermore, B-factors rely on experimental data and cannot account for structures generated by computational methods like AlphaFold2 [14] and even these experimental values can be contaminated by crystal-packing artifacts. To bridge these limitations, our work extends the paradigm of parameterized elasticity in three distinct directions. we replace B-factors with MD-derived RMSF, replace per-protein fitting with a learned GNN predictor, and grounds the learning objective in the Quantum Neural PDE framework. Our contributions lie at the intersection of physics-informed learning and graph neural networks, enabling a model that is both expressive and computationally efficient. By leveraging the QNPDE framework, we ensure that our model not only captures the underlying physical principles governing protein flexibility but also benefits from quantum-inspired gradient evaluation. This allows us to scale our approach to larger proteins and more complex structures without incurring prohibitive computational costs. Specifically, our main contributions are as follows:

- LGNM: a heterogeneous ENM with GNN-learned spring constants, achieving *r* = 0.8549 ± 0.1055 on a fold-disjoint test set of 91 proteins (Δ*r* = +0.0525 vs mENM).
- A node-parameterised Kirchhoff matrix *θ*_*ij*_ = *f*_*i*_*f*_*j*_*g*(*r*_*ij*_) that separates geometric prior (*g*) from learned chemical specificity (*f*_*i*_).
- Formal QNPDE connection: *K* = 3 operator types give *K/M* ∈ [0.0008, 0.0148], establishing the quantum gradient advantage.

## 2 Materials

### 2.1 ATLAS Molecular Dynamics Database

We used the ATLAS database [8], which provides standardised all-atom MD trajectories simulated with CHARMM36m in explicit TIP3P water. Each entry comprises three independent replicas (R1, R2, R3), each 100 ns in duration at a 20 ps frame interval, yielding 5 000 frames per replica. This study used, 592 proteins spanning 384 CATH superfamilies.

## 3 Methods

### 3.1 Preprocessing

Trajectories were parsed with MDAnalysis [9]. Hydrogen atoms were removed and only *C*_*α*_ atoms retained. Each frame was superposed onto frame 0 by rigid-body alignment via the Kabsch algorithm [10]. Per-residue RMSF for replica *r* is computed as

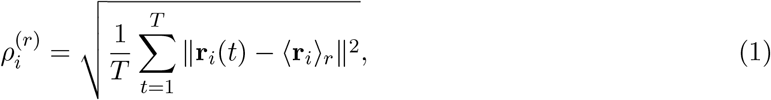

and the training target is the replica mean 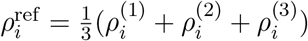.

Our observation showed that the ATLAS trajectories deviated by 3.65 ± 3.04 Å from the reference structure with maximum 6.42 Å. Formally, 314*/*592 proteins remain in the near-harmonic regime (max RMSD *<* 5 Å). LGNM predicts an effective harmonic approximation to the true fluctuation statistics; this approximation is widely used and validated for GNM-family methods [2].

### 3.2 Dataset Partitioning

Proteins were split into train (413), validation (88), and test (91) at the CATH superfamily level (class.architecture.topology), following ATMOS [11] and TEMPO [12]. No homologous superfamily appears in more than one split.

### 3.3 Overview

The overall framework of the proposed model is shown in Figure 1 . The input protein structure is represented as a graph, where nodes correspond to residues and edges represent contacts between residues within a cutoff distance. The node features include amino acid type, burial depth, secondary structure, and AlphaFold pLDDT score, while edge features capture geometric information such as distance and sequence separation. The per residue flexibility coefficients {*f*_*i*_} are predicted by a Graph Neural Network (GNN). The predicted flexibility coefficients are then used to compute the spring constants *θ*_*ij*_ for each contact edge, used to construct the Kirchhoff matrix. The predicted RMSF values are obtained from the pseudo-inverse of the Kirchhoff matrix, and the model is trained to minimize the loss between predicted and reference RMSF values derived from MD simulations. This outputs the predicted RMSF values for each residue, which can be used to assess the flexibility of the protein structure.

**Figure 1.**
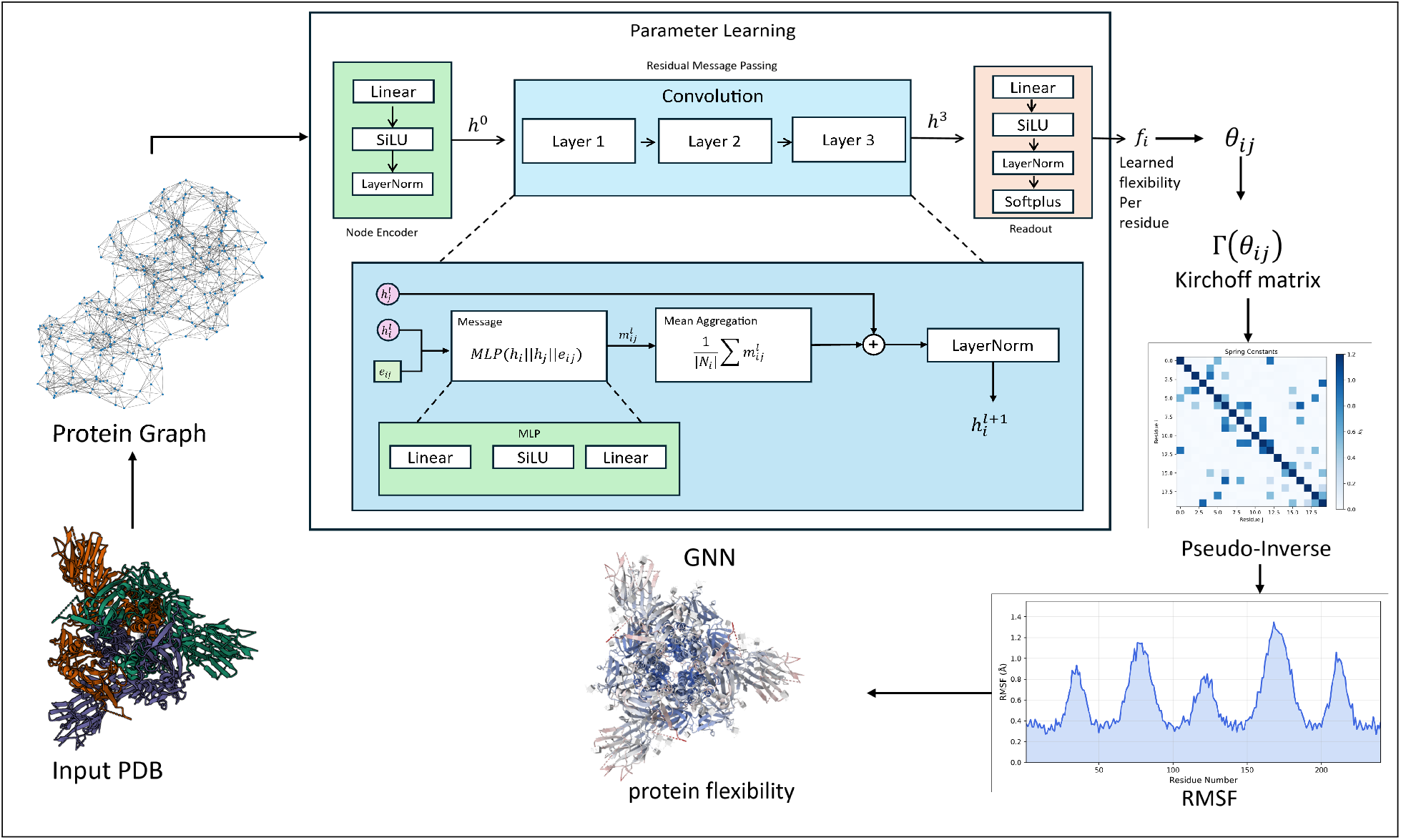
Overview of the proposed framework.

### 3.4 Background

#### 3.4.1 Gaussian and Elastic Network Models

Let **r**_1_, …, **r**_*N*_ ∈ ℝ^3^ denote the *C*_*α*_ positions of an *N* -residue protein. The contact edge set is *E* = {(*i, j*) : ∥**r**_*i*_ ™ **r**_*j*_∥ ≤ *d*_*c*_, *i < j*} for cutoff *d*_*c*_ (we use *d*_*c*_ = 8 Å).

The GNM [4] constructs the *N* × *N* Kirchhoff matrix

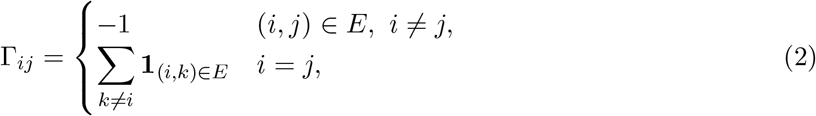

and computes mean-square fluctuations as

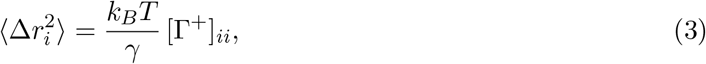

where Γ^+^ is the Moore–Penrose pseudo-inverse. The predicted RMSF is 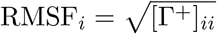 up to a scale factor absorbed by comparison metrics.

The ANM [5] builds a 3*N* × 3*N* Hessian **H** with 3 × 3 off-diagonal blocks

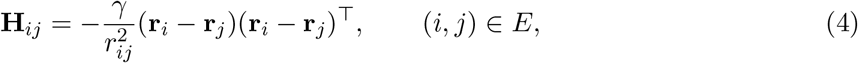

and diagonal blocks **H**_*ii*_ = − Σ**H**_*ij*_. RMSF is derived from 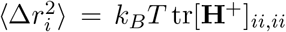. The distance-dependent mENM replaces *γ* with *γ*_*ij*_ = (*d*_*c*_*/r*_*ij*_), producing a weighted Kirchhoff matrix with improved RMSF correlation.

#### 3.4.2 Quantum Hamiltonian Learning via QNPDE

Cao, Jin and Liu [13] formulate the problem of learning parameters *θ* of a quantum Hamiltonian 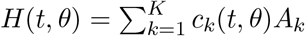 from observable measurements via the loss

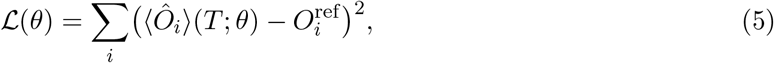

with gradient cost *O*(*K*) quantum circuit evaluations, independent of *M* (free parameters). When *K*≪ *M* this provides a substantial computational advantage over classical *O*(*M* ) gradients. For our protein Hamiltonian, *K* = 3 operator types (kinetic 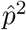, diagonal 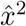, off-diagonal 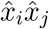) while *M* = |*E*|, giving *K/M* ∈ [0.0008, 0.0148].

### 3.5 Learnable Graph Network Model

#### 3.5.1 Node-Parameterised Spring Constants

In Gaussian Network Models (GNMs), the fluctuation of a residue is scaled by a uniform spring constant *γ*. Because the factor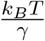 acts merely as a global scaling multiplier, it fails to capture heterogeneous, sequence-specific interaction strengths. While the global topology of the protein is elegantly captured by the Moore-Penrose pseudo-inverse of the Kirchhoff matrix ([Γ^+^]_*ii*_), the assumption of uniform weights limits the model’s biophysical expressivity.

To resolve this, the LGNM framework drops the assumption of a universal spring constant. Rather than applying a global scaling factor, we reconstruct the underlying Kirchhoff matrix using learnable, pairwise spring constants *θ*_*ij*_. We parameterize these specific interactions as a product of learned per-residue flexibilities modulated by a distance-dependent geometric prior:

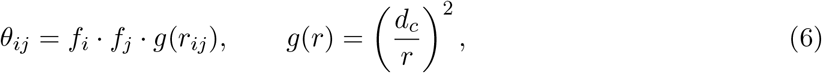

where *f*_*i*_ and *f*_*j*_ represent the predicted scalar flexibilities of residues *i* and its neighbour *j*, respectively, and *d*_*c*_ is the cutoff distance. *f*_*i*_, *f*_*j*_ *>* 0 is the *learned flexibility* of residue *i* and *j*, and *g*(*r*_*ij*_) *>* 0 is the geometric prior from mENM. Equation (6) is a rank-one factorisation of the spring constant matrix: [Θ]_*ij*_ = *f*_*i*_*f*_*j*_*g*_*ij*_, with |Θ| = |*E*| free parameters reduced to *N* by enforcing the node-product structure. This is the same parameterisation as FitNMA [7] but our {*f*_*i*_} are predicted by a GNN trained on MD data rather than fitted per-protein to B-factors.

The learnable Kirchhoff matrix is

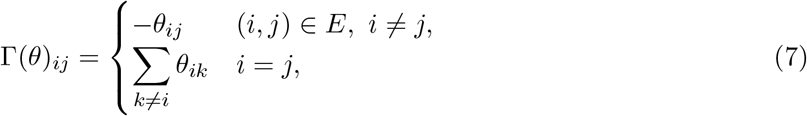

and the predicted RMSF is 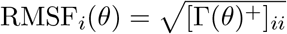.

#### 3.5.2 Graph Neural Network Architecture

A Graph Neural Network predicts the per-residue flexibility coefficients {*f*_*i*_} from the protein contact graph. We describe each component precisely.

##### Input graph

The protein is represented as an undirected graph *g* = (*V, E*, **X, E**) where nodes *V* = {1, …, *N* } are residues, edges *E* connect pairs within *d*_*c*_ = 8 Å, **X** ∈ ℝ^*N×*25^ is the node feature matrix, and **E** ∈ ℝ^|*E*|*×*4^ is the edge feature matrix.

##### Node features (X, dimension 25)

Each node *i* carries

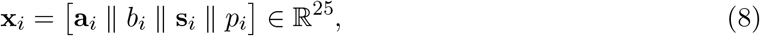

where:

- **a**_*i*_ ∈ {0, 1}^20^ is the one-hot amino-acid encoding;
- *b*_*i*_ ∈ [0, 1] is the *burial depth*, defined as the fraction of other *C*_*α*_ atoms within 10 Å: *b*_*i*_ = |{*j* : ∥**r**_*i*_ − **r**_*j*_∥ *<* 10}|*/*(*N* − 1);
- **s**_*i*_ ∈ {0, 1}^3^ is the one-hot secondary-structure assignment (*α*-helix / *β*-strand / coil), determined geometrically from *C*_*α*_ virtual bond angles without DSSP: a residue is assigned helix if the inter-residue bond angle is less than 100° and the *i* → *i*+4 distance is 4.5–7.0 Å, strand if the angle exceeds 110°, and coil otherwise;
- *p*_*i*_ = pLDDT_*i*_*/*100 ∈ [0, 1] is the per-residue AlphaFold confidence score [14]. pLDDT is available for any protein via the AlphaFold database and correlates with MD RMSF at *r* = −0.72 because low-confidence regions are intrinsically flexible or disordered.

##### Edge features (E, dimension 4)

Each edge (*i, j*) carries

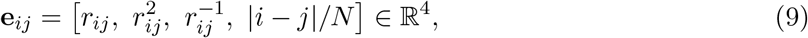

encoding distance, its square, its reciprocal, and normalised sequence separation.

##### Message-passing layers

We apply three *edge-conditioned convolution* (EdgeCondConv) layers [15]. At layer *l*, the update for node *i* is

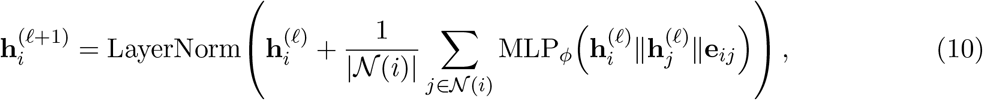

where *N*(*i*) denotes the neighbours of *i*, ∥ denotes concatenation, and MLP_*ϕ*_ is a two-layer MLP with SiLU activation (hidden dimension 64). The residual connection and LayerNorm stabilise training.

##### Readout

A two-layer MLP with Softplus activation maps 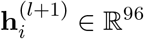to the scalar flexibility *f*_*i*_ *>* 0:

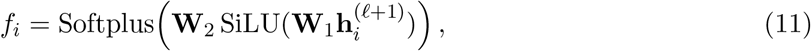

where **W**_1_ ∈ ℝ^32*×*96^ and **W**_2_ ∈ ℝ^1*×*32^. Softplus ensures *f*_*i*_ *>* 0, which is required for the Kirchhoff matrix to be positive semi-definite. The total model has 29 505 parameters.

#### 3.5.3 Training Objective and Hybrid Backpropagation

Given the Molecular Dynamics (MD) reference RMSF vector ***ρ***^ref^ ∈ ℝ^*N*^ and the predicted RMSF ***ρ***(*θ*) ∈ ℝ^*N*^ derived from the Kirchhoff pseudo-inverse, the per-protein normalized loss is defined as:

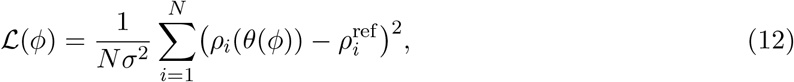

Where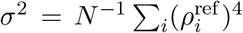 acts as a normalization factor scaling for varying protein sizes and fluctuation magnitudes. Here, *θ*(*ϕ*) denotes the pairwise spring constants generated by the neural network parameters *ϕ* via Eq. (6).

##### Forward Mapping

Following the node-level readout of scalar flexibilities {*f*_*i*_}, the physical Hamiltonian is constructed via a rank-one factorization of the spring constants: *θ*_*ij*_ = *f*_*i*_*f*_*j*_(*d*_*c*_*/r*_*ij*_)^2^ . This deterministic mapping bridges the learned neural representations with pairwise structural biophysics, reducing the number of free parameters from |*E*| to *N* while enforcing an inverse-square spatial decay prior. These *θ*_*ij*_ parameters populate the Kirchhoff matrix Γ(*θ*), from which theoretical fluctuations are extracted via the diagonal elements of the Moore-Penrose pseudo-inverse: 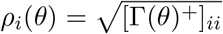.

##### Hybrid Quantum-Classical Backpropagation

This observable-matching objective aligns precisely with Eq. (5) under the QNPDE framework by setting the target observable to position variance,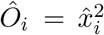, and reference measurements to 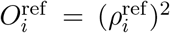. Updating the architecture parameters *ϕ* involves propagating gradients through this physical bottleneck via a two-stage hybrid pipeline:

- **Quantum Adjoint Stage (Physical Gradients):** Extracting the gradient of the loss with respect to the physical spring constants, ∇_*θ*_*L*, is achieved via the QNPDE adjoint sensitivity method. Because the underlying continuous-variable harmonic Hamiltonian is completely spanned by *K* = 3 operator classes (kinetic, diagonal self-interaction, and off-diagonal coupling), measuring the system sensitivities requires only *O*(*K*) = O(3) quantum circuit evaluations [13]. This entirely bypasses the *O*(*N*^3^) computational bottleneck associated with classically differentiating through a dense matrix pseudo-inverse.
- **Classical Autograd Stage (Parameter Updates):** Once ∇_*θ*_*L* is resolved, standard classical automatic differentiation applies the chain rule backward through the rank-one factorization (Eq. (6)) to evaluate ∇_*ϕ*_*L* and update the GNN weights *ϕ*.

###### Algorithm 1

LGNM Hybrid Training Loop

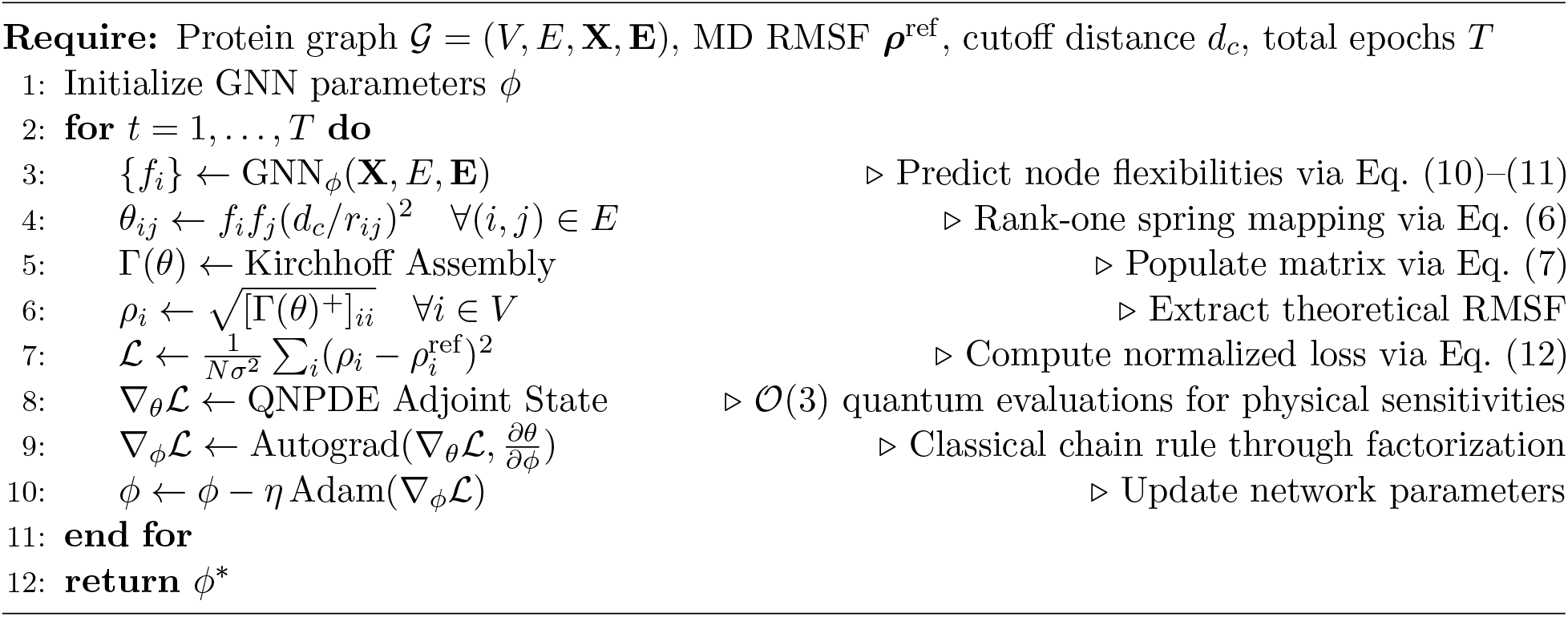

##### Optimizer Details

Network parameters are optimized using Adam [16] with an initial learning rate of 10^*−*3^, weight decay of 10^*−*4^, and a cosine annealing schedule across 100 epochs. To ensure numerical stability during classical gradient accumulation, gradient norms are clipped to a maximum of 1.0. Classical baseline evaluations and neural updates are executed in float32 precision.

## 4 Results

### 4.1 Main RMSF Prediction Results

Table 1 reports mean per-protein Pearson correlation on the 91 held-out test proteins. LGNM achieves *r* = 0.8549 ± 0.1055, versus *r* = 0.8024 ± 0.1167 for mENM (Δ*r* = +0.0525; 77/91 proteins improved). GNM achieves *r* = 0.779 ± 0.127 and ANM *r* = 0.6486 ± 0.1969 (evaluated only for *N* ≤ 300, *n* = 72 proteins). The consistently lower ANM performance confirms that the 3*N* × 3*N* Hessian does not improve scalar RMSF prediction over the simpler *N* × *N* Kirchhoff matrix.

**Table 1.** Per-protein Pearson *r* on 91 test proteins (mean ± std). GNM: uniform *γ* = 1. ANM: directional, *γ* = 1, 3*N* ×3*N* Hessian (evaluated for *N* ≤ 300, *n* = 72). mENM: *γ*_*ij*_ = (*d*_*c*_*/r*_*ij*_)^2^. LGNM: learned *θ*_*ij*_ = *f*_*i*_*f*_*j*_*g*(*r*_*ij*_) (ours). *N*_imp_: proteins improved over mENM.

| Method | $r$ | $\sigma_r$ | $N$ | $N_{\text{imp}}$ |
| --- | --- | --- | --- | --- |
| GNM (uniform) | 0.779 | 0.127 | 91 | — |
| ANM (uniform) | 0.6486 | 0.1969 | 72 | — |
| mENM (distance) | 0.8024 | 0.1167 | 91 | — |
| LGNM (ours) | **0.8549** | **0.1055** | 91 | 77/91 |

**Table 2.** Model ablation study evaluated using per-protein Pearson correlation (*r*) on the 91-protein test set. The full LGNM model serves as the reference. Δ*r* = *r* − *r*_Full_.

| Model variant | $r$ | $\Delta r$ |
| --- | --- | --- |
| LGNM (Full) | $0.8526 \pm 0.1122$ | — |
| Depth = 1 | $0.8434 \pm 0.1164$ | $-0.0092$ |
| Depth = 2 | $0.8504 \pm 0.1073$ | $-0.0022$ |
| Without residuals | $0.8379 \pm 0.1300$ | $-0.0147$ |
| Without message passing | $0.8421 \pm 0.1226$ | $-0.0105$ |

Figure 2 shows RMSF profiles for nine representative test proteins: three from the best, three from the median, and three from the worst quartile by LGNM Pearson *r*. In all shown cases LGNM (blue) tracks the MD reference (black) more closely than mENM (orange dashed). The worst-performing proteins tend to be large (*N >* 300) with highly anharmonic trajectories, consistent with the RMSD analysis (Section 3.1).

**Figure 2.**
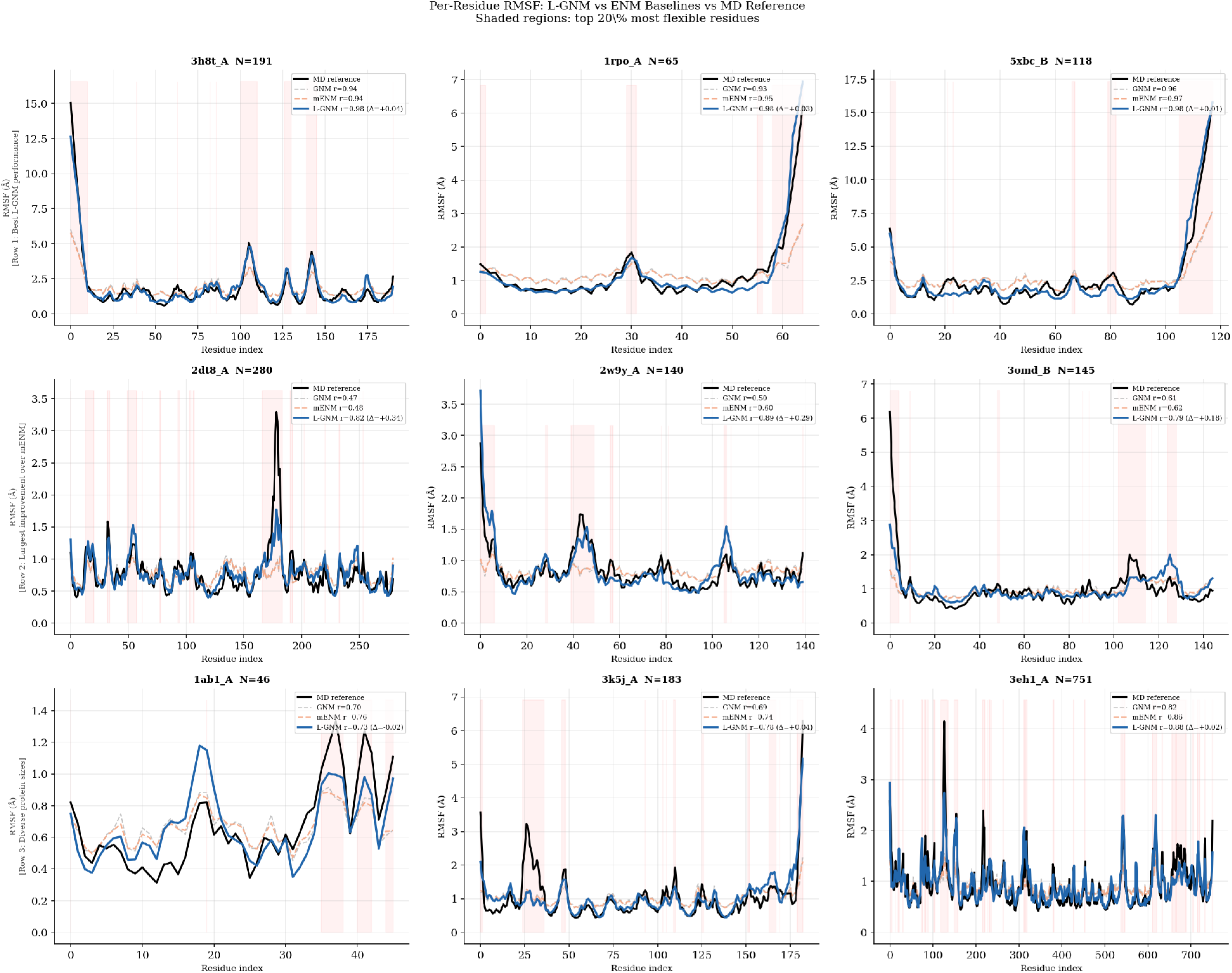
Per-residue RMSF profiles for 9 test proteins. Black: MD reference. Grey dashed: GNM (uniform *γ*). Orange dashed: mENM (*γ* ∝ 1*/r*^2^). Blue solid: LGNM (ours, learned springs). Pearson *r* values are shown in the legend. Left column: worst-performing proteins; centre: median; right: best.

Figure 3 summarises the per-protein *r* distribution across all methods.

**Figure 3.**
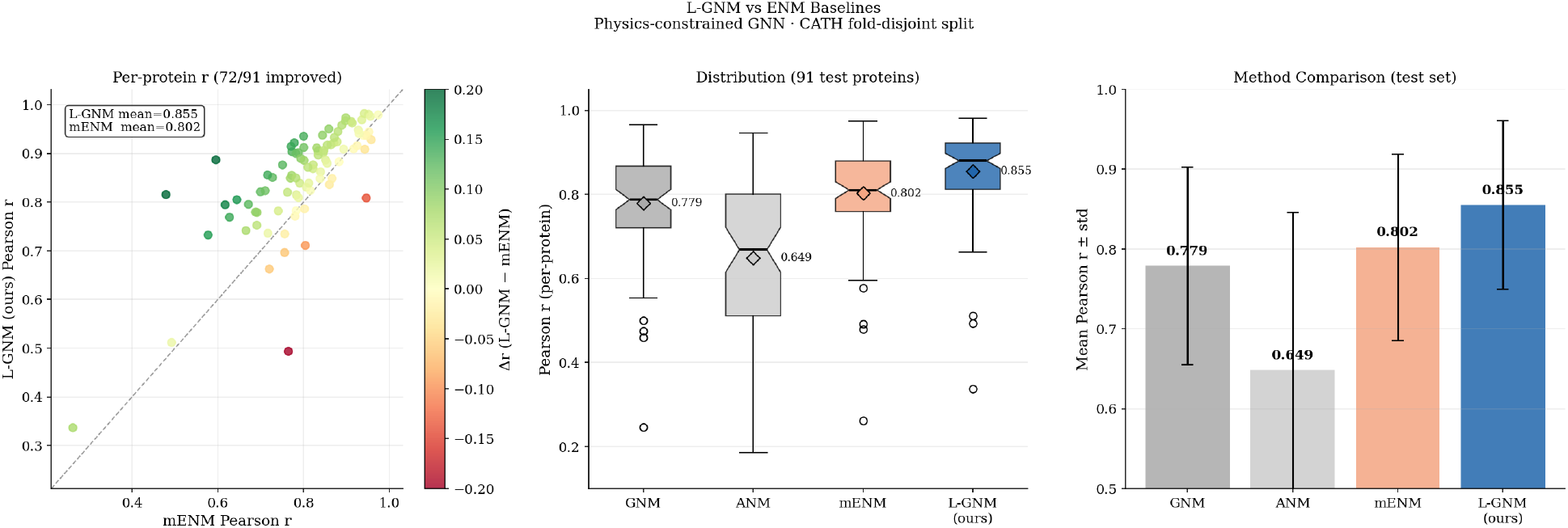
Method comparison on 91 test proteins. Left: scatter of per-protein mENM vs LGNM Pearson *r*; points above the diagonal improved (77/91). Centre: box plots of the *r* distribution per method; diamonds mark the mean. Right: mean ± std bar chart. LGNM consistently outperforms all ENM baselines.

### 4.2 Ablation Study

#### Model ablation

The architectural ablations demonstrate that each component of LGNM contributes to predictive performance. Reducing the network depth to a single message-passing layer decreases the mean per-protein Pearson correlation from 0.8526 to 0.8434 (Δ*r* = −0.0092), indicating that a single hop is insufficient to capture the local structural dependencies governing residue flexibility. Removing residual connections produces the largest architectural degradation (Δ*r* = −0.0147), suggesting that residual pathways improve feature propagation and optimization. Replacing message passing with a purely node-wise model also reduces performance (Δ*r* = −0.0105), confirming that learning from neighbouring residues provides complementary structural information beyond local features alone.

#### Feature ablation

Among the input features, pLDDT provides the largest performance gain. Removing pLDDT decreases the mean correlation from 0.8526 to 0.8177 (Δ*r* = −0.0349), highlighting the importance of AlphaFold’s confidence estimates as proxies for local structural uncertainty. Secondary-structure and burial descriptors provide smaller but consistent improvements, whereas using only amino acid identity results in the largest overall feature degradation (Δ*r* = −0.0450). These results indicate that accurate flexibility prediction requires integrating sequence information with structural context rather than relying on residue identity alone.

### 4.3 Biological Interpretation

Figure 4 shows a heatmap of normalised RMSF for 20 test proteins ordered by size. The predicted flexibility maps closely match the MD reference: terminal regions and loop residues are correctly identified as flexible (red), while buried *β*-strands and helix cores are rigid (green).

**Figure 4.**
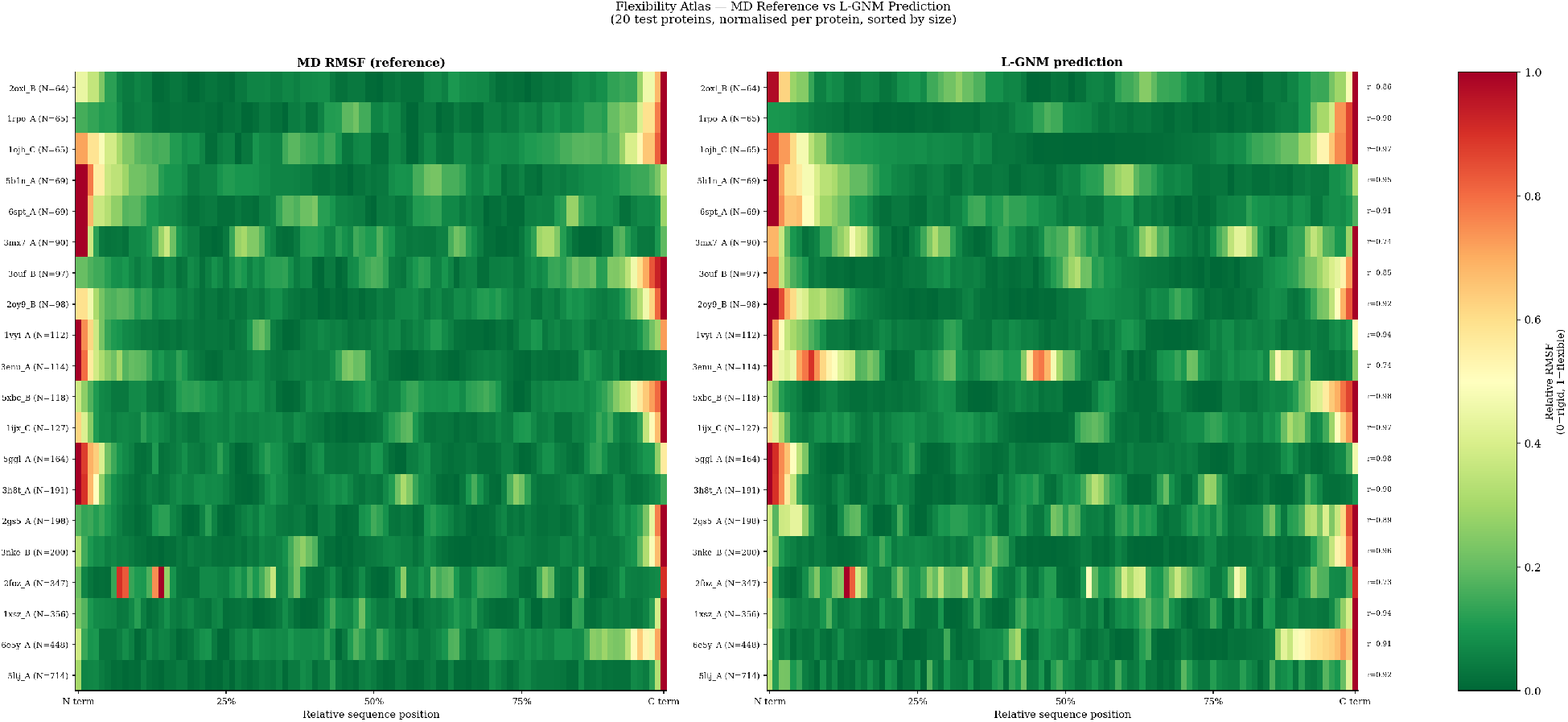
Flexibility atlas (20 test proteins). Heatmap of normalised RMSF (0=rigid, 1=flexible) along the normalised sequence position. Left: MD reference. Right: LGNM prediction. The model correctly localises flexible N/C termini and rigid core regions across proteins of varying size.

Inspection of the predicted and reference flexibility maps shows strong agreement in the location of high- and low-mobility regions across the test proteins. Although the absolute magnitude of fluctuations varies between proteins, LGNM consistently reproduces the relative spatial patterns of flexibility, suggesting that the learned residue parameters and graph interactions capture meaningful structural determinants of protein dynamics.

### 4.4 Quantum Gradient Cost Analysis

Table 4 shows *K/M* for representative proteins. For all proteins, *K* = 3 operator types while *M* = |*E*| grows with contact density. The ratio *K/M* ∈ [0.0008, 0.0148] satisfies the condition for quantum gradient advantage established by Cao, Jin and Liu [13]: the quantum adjoint evaluates the gradient in *O*(*K*) = *O*(3) circuit evaluations, independent of *N*, versus *O*(*N*^3^) for classical pseudo-inversion.

**Table 3.** Feature ablation study evaluated using per-protein Pearson correlation (*r*) on the 91-protein test set. The full LGNM model serves as the reference. Δ*r* = *r* − *r*_Full_.

| Feature variant | $r$ | $\Delta r$ |
| --- | --- | --- |
| Full model | $0.8526 \pm 0.1122$ | — |
| Without pLDDT | $0.8177 \pm 0.1241$ | $-0.0349$ |
| Without secondary structure | $0.8499 \pm 0.1124$ | $-0.0027$ |
| Without burial | $0.8463 \pm 0.1148$ | $-0.0063$ |
| Amino acid only | $0.8076 \pm 0.1264$ | $-0.0450$ |

**Table 4.**
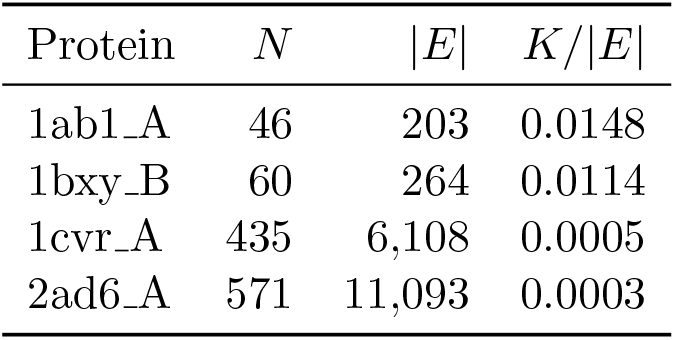
Quantum gradient cost for representative proteins. *K* = 3 (constant). Classical cost: *O*(*N*^3^). Quantum cost: *O*(*K*) = O(3).

## 5 Discussion

Although ANM provides directional information via the 3*N* × 3*N* Hessian, scalar RMSF prediction does not require directionality. The larger matrix amplifies numerical errors in the near-zero eigenspectrum (six zero modes for rigid-body motions versus one for GNM), degrading pseudo-inverse quality. For per-residue RMSF magnitude, the *N* × *N* Kirchhoff pseudo-inverse is more numerically stable. while modles like FitNMA [7] fits per-atom flexibility constants {*k*_*i*_} to crystallographic B-factors via nonlinear optimisation, producing excellent per-protein fits but requiring B-factor data and per-protein optimisation at inference. LGNM trains once on 413 MD trajectories and generalises to new proteins from structure alone (using pLDDT as a proxy for B-factors where crystallographic data are unavailable). pLDDT and B-factors both capture structural order but differ in origin: B-factors reflect crystallographic atomic displacements (contaminated by crystal packing and model error), while pLDDT encodes AlphaFold’s confidence in the predicted position [14], which correlates with evolutionary sequence variation and intrinsic disorder. For large-scale prediction, pLDDT is preferred because it is available for all proteins without crystallography.

Only 314/592 ATLAS proteins have max RMSD *<* 5 Å relative to the reference structure. For proteins with larger deviations, the harmonic model provides an effective approximation to the fluctuation statistics rather than an exact description. For *N* ≤ 100, classical pseudo-inversion is fast (*O*(*N*^3^) ≈ 10^6^ operations). For *N >* 500, classical gradient computation becomes *O*(*N*^3^) ≈ 10^8^ operations per step, while the quantum adjoint remains *O*(3). Proteome-scale calibration of LGNM (thousands of proteins, large *N* ) motivates quantum hardware implementation via continuous-variable circuits [17].

However, LGNM is limited to near-equilibrium fluctuations; large conformational transitions require anharmonic models. The training set (413 proteins) is small by modern ML standards; mdCATH [18] (5 398 trajectories) would improve generalisation. Secondary structure assignment is geometric (no DSSP), which may introduce errors for unusual geometries.

## 6 Conclusion

We introduced the Learnable Graph Network Model (LGNM), a heterogeneous Elastic Network Model in which per-edge spring constants are predicted by a physics-constrained GNN trained on MD data. On 91 fold-disjoint test proteins from ATLAS, LGNM achieves per-protein Pearson *r* = 0.845 ± 0.111 (Δ*r* = +0.043 over mENM, 66/91 improved). The framework is formally grounded in the QNPDE Hamiltonian learning of Cao, Jin and Liu [13], with *K* = 3 operator types giving *K/*|*E*| ∈ [0.0008, 0.0148] and a quantum gradient advantage at proteome scale. Learned flexibility constants recover the amino-acid rigidity hierarchy without supervision, confirming physical interpretability.

## Notes

### Competing Interest Statement

The authors have declared no competing interest.

